# NQO1 binds and supports SIRT1 function

**DOI:** 10.1101/139907

**Authors:** Peter Tsvetkov, Julia Adler, Yaarit Adamovich, Gad Asher, Nina Reuven, Yosef Shaul

## Abstract

Silent information regulator 2-related enzyme 1 (SIRT1) is an NAD^+^-dependent class III deacetylase and a key component of the cellular metabolic sensing pathway. The requirement of NAD^+^ for SIRT1 activity led us to assume that NQO1, an NADH oxidoreductase producing NAD^+^, regulates SIRT1 activity. We show here that SIRT1 is capable of increasing NQO1 (NAD(P)H Dehydrogenase Quinone 1) transcription and protein levels. NQO1 physically interacts with SIRT1 but not with an enzymatically dead SIRT1 H363Y mutant. The interaction of NQO1 with SIRT1 is markedly increased under mitochondrial inhibition. Interestingly, under this condition the nuclear pool of NQO1 is elevated. Depletion of NQO1 compromises the role of SIRT1 in inducing transcription of several target genes and eliminates the protective role of SIRT1 following mitochondrial inhibition. Our results suggest that SIRT1 and NQO1 form a regulatory loop where SIRT1 regulates NQO1 expression and NQO1 binds and mediates the protective role of SIRT1 during mitochondrial stress. The interplay between an NADH oxidoreductase enzyme and an NAD^+^ dependent deacetylase may act as a rheostat in sensing mitochondrial stress.

---

SIRT1 is a member of the NAD^+^ dependent class III histone/protein deacetylase family that is composed of seven members (SIRT1-7) that are evolutionarily conserved (1). The Sirtuins are differentially localized in the cell. SIRT3, SIRT4, and SIRT5 are mitochondrial, SIRT2 is cytoplasmic whereas SIRT6, SIRT7 and SIRT1 are mainly nuclear (2). SIRT1, the yeast Sir2 ortholog, is the most studied sirtuin family member. SIRT1 is active in the deacetylation of histones, transcription factors and enzymes involved in key cellular processes such as metabolism (3,4), longevity (5), circadian oscillation (6) and differentiation (7). Moreover, NAD^+^ plays a crucial role in activating SIRT1 activity. Under caloric restriction or fasting, the level of NAD^+^ rises and SIRT1 function is induced (4,8,9). Increased levels of NAD^+^ due to elevated NAD^+^ salvage pathway synthesis, external NAD^+^ precursor addition or the inhibition of NAD^+^ utilizing enzymes such as PARP all have been shown to promote SIRT1 activity (10-12).

NQO1 is a ubiquitous flavoenzyme that catalyzes two-electron reduction of various quinones, utilizing NADH as an electron donor and generating NAD^+^ (13). In recent years it has become evident that NQO1 has a dual function in the cell. First, by associating with the 20S proteasome, it regulates the degradation of several intrinsically disordered proteins (IDPs) (as summarized in (14)). The second and more studied NQO1 role is as part of the cellular defense mechanism in response to electrophilic and/or oxidative stress generated by exposure to chemicals or endogenous quinones (15). NQO1 enzymatic activity shifts the cellular redox state as was shown in NQO1 knockout mice that exhibited a significant increase in the NADH/NAD^+^ ratio (16). This is also the case in the involvement of NQO1 in the plasma membrane electron transfer (PMET) system highly regulating the NADH/NAD^+^ ratio and cellular viability following different cellular stresses, including mitochondrial stress (17).

Previously we proposed that the role of NQO1 in regulating the NADH/NAD+ ratio might regulate SIRT1 activity (18). Recently we reported that NQO1 regulates the protein levels of transcription coactivator PGC-1α, a key metabolic gene regulator (19). The stabilizing effect of NQO1 on PGC-1α is NADH-dependent and regulates PGC-1α accumulation and transcriptional activity following cellular starvation (19). PGC-1α is also activated by deacetylation by SIRT1 that in turn is regulated by the alteration in the cellular NADH/NAD+ ratio (4). These findings and the ability of NQO1 to alter the NADH/NAD+ ratio in cells led us to explore a possible cross talk between SIRT1 and NQO1. Here we report on a functional physical interaction between NQO1 and SIRT1. We further demonstrate that NQO1 positively regulates SIRT1 activity in response to mitochondrial stress.

## EXPERIMENTAL PROCEDURES

### Cells and Cell culture

293 human kidney cells (HEK) and NIH3T3 cells were grown in Dulbecco’s modified Eagle’s medium (DMEM) supplemented with 10% fetal bovine serum (FBS), 100 units/ml penicillin, and 100 mg/ml streptomycin and cultured at 37°C in a humidified incubator with 5.6% CO2. C2C12 cells were grown as previously described (19). H1299 with a YFP insertion in NQO1 exon 2 (clone 130207p11G9) was obtained from the laboratory collection of Uri Alon (http://www.weizmann.ac.il/mcb/UriAlon/DynamProt/index.html).

### Compounds

Resveratrol, rotenone, antimycin A, oligomycin, nicotinamide and cycloheximide were purchased from Sigma. EX527 was from PeproTech.

### Plasmids, transfection and retroviral infection

The plasmids used were: pEFIRES NQO1, NQO1 C609T, flag-NQO1, flag-p73β. pEFIRES flag-SIRT1 and flag-SIRT1 H363Y, pcDNA3.1 flag-SIRT2, pcDNA3.1 flag-SIRT3 (provided by Eric Verdin), pBabe SIRT1 and pBabe SIRT1 H363Y (provided by Robert Weinberg), pcDNA3.1 flag-SIRT4-7 (provided by Haim Cohen) and the pCDNA3 Peredox-mCherry reporter (provided by Gary Yellen). Transient transfections of HEK293 cells were carried out by the calcium phosphate method. For retroviral infection HEK293T cells were transfected with pBabe GFP, pBabe SIRT1 or SIRT1 H363Y mutant together with Ψ-helper plasmid. Two days after transfection the supernatants containing the viral particles were harvested and used to infect NIH3T3 and C2C12 cells. NQO1 knockdown was achieved in C2C12 by introducing pLKO. 1 shRNA against NQO1 (Sigma) and nontargeting control shRNA (Sigma) with lentivirus transduction. Lentiviral particles were produced in HEK293T cells according to the manufacturer’s protocol.

### Immunoblot Analysis

Cell extracts and immunoblot analysis were carried out as previously described (20). The antibodies used were: goat anti NQO1 C19 and R20 (Santa Cruz), rabbit anti SIRT1 H300 (Santa Cruz) and mouse monoclonal anti His (Sigma).

### Co-immunoprecipitation Studies

Co-immunoprecipitation experiments were carried out as previously described (20). Briefly, extracts from cells transiently transfected with indicated plasmids were incubated with Flag beads (Sigma), washed, eluted and separated on a 12.5% SDS-PAGE. *In vitro* coimmunoprecipitation experiment with recombinant purified His-SIRT1 and NQO1 was carried out with Ni-NTA beads (Qiagen) in buffer containing 50 mM Tris-HCl pH 8.0, 150 mM NaCl and 20 mM imidazole. Following washes samples were eluted in the same buffer with 200 mM imidazole. *Purification of recombinant proteins*. pET28-His-TEV-NQO1 and pET28-His-SIRT1 were expressed in bacteria and cells were lysed by sonication in 50 mM Tris-HCl pH 7.5, 150 mM NaCl, Lysozyme and 1 mM PMSF. Soluble His-TEV-NQO1 and His-SIRRT1 were purified using Ni-NTA column (HiTrap chelating HP) followed by gel filtration chromatography (HiLoad 16/60 superdex 200). Purified His-TEV-NQO1 was further cleaved by TEV protease and the His-TEV was removed upon binding to Ni-NTA column. For co-immunoprecipitation of endogenous SIRT and NQO1, mouse livers RIPA extracts were immunoprecipitated with either control protein A/G beads (Santa Cruz) or A/G beads coated with either non-immune rabbit IgG, or rabbit anti SIRT1 H300 (Santa Cruz). Protein levels were analyzed by Western blotting.

### Immunofluorescence analysis

Cells were fixed with 4% paraformaldehyde, permeabilized with 0.5% Triton-X 100 and blocked with fetal calf serum containing 10% (v/v) skim milk. Cells were then incubated with goat anti NQO1 C19 and R20 antibodies (Santa Cruz). Following incubation with Cy5-conjugated donkey anti-goat antibody (Jackson Immuno Reasearch Laboratories), coverslips were mounted in DAPI Fluoromount-G (SouthernBiotech, Birmingham, AL, USA). Microscopic images were obtained using laser scanning microscope LSM710 (Carl Zeiss, Microimaging GmbH, Göttingen, Germany) with plan-apochromat 63 × /1.40 oil DIC M27 objective, and managed by Laser Sharp 2000 software (Zeiss, Munich, Germany). Representative images with identical laser intensities.

### NQO1 activity assay

Cells were plated in 96 well plates and treated as described in the figure legend. NQO1 activity was assayed as previously described (21).

### Viability assay

Cell viability was analyzed using the XTT assay (Biological Industries) and spectrophotometrically quantified according to manufacturer’s instructions.

### Pulse-chase experiment

Pulse-chase experiments were conducted as described previously with slight modifications (22). HEK293T cells were incubated with medium depleted of methionine (−M) for 20 min. [^35^S]Methionine (10 mCi/ml; Amersham Biosciences) was added reaching a final concentration of 0.2 mCi/μl for a pulse of 30 min. Cells were washed, and medium containing 2% unlabeled methionine was added for the indicated time (chase). Cells were collected, and protein extracts were subjected to immunoprecipitation as described above.

### Gene expression analysis

Total RNA was isolated from cells by using Tri reagent (Molecular Research Center), and reverse transcribed using the iScript cDNA kit (Bio-Rad). Gene expression levels were normalized to the geometric mean of three housekeeping genes, TATA binding protein (TBP), hypoxanthine phosphoribosyltransferase (HPRT), and glyceraldehyde-3-phosphate dehydrogenase (GAPDH) and expressed relative to control using the threshold cycle (ΔΔ*C_T_*) method (19). The sequences of the primers used in this study will be provided upon request.

## RESULTS

### NQO1 binds SIRT1

To investigate NQO1 interaction with the Sirtuin family members we performed co-immunoprecipitation assays. Flag SIRT1-7 and p73β as a negative control were transfected in the presence of NQO1 in HEK293 cells. The Sirtuins were immunoprecipitated with flag beads and the levels of co-immunoprecipitated NQO1 was analyzed. NQO1 showed high association with SIRT1 (Fig. 1A). Mild association of NQO1 was also observed with SIRT2, SIRT6 and SIRT7. Next we asked whether SIRT1 functional conformation is required for NQO1 binding. SIRT1 H363Y is a catalytically inactive form of SIRT1. The SIRT1 mutant did not bind NQO1 (Fig. 1B) suggesting a functional conformation of SIRT1 is required for the interaction with NQO1. To further validate the direct physical association between SIRT1 and NQO1 we bacterially expressed and purified both proteins. NQO1 was incubated alone or in the presence of His-SIRT1 followed by precipitation with Ni-beads. NQO1 precipitated only in the presence of His-SIRT1 (Fig. 1C) suggesting the binding between SIRT1 and NQO1 is direct.

**Figure 1.**
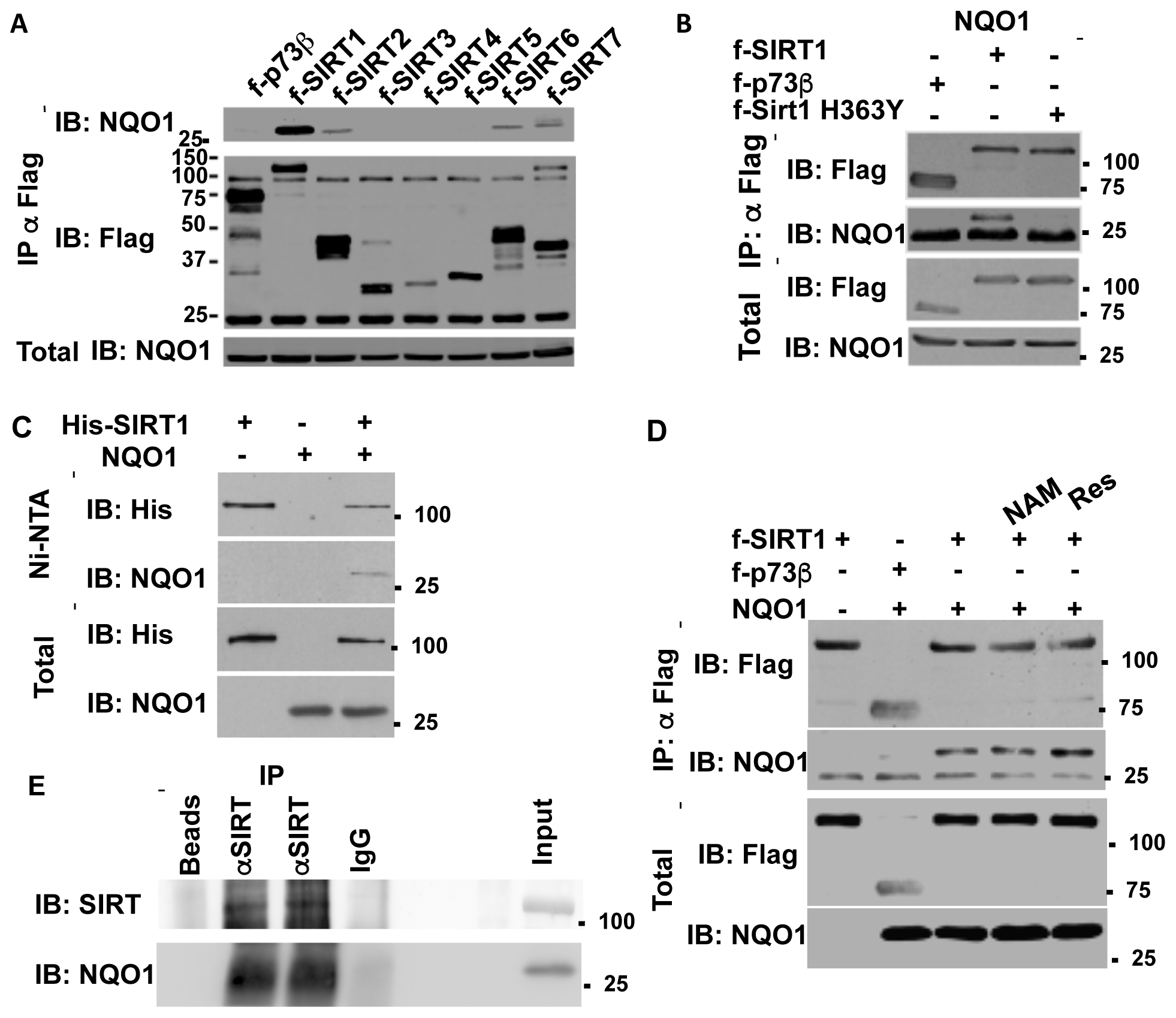
SIRT1 directly binds NQO1. A) HEK293 cells were co-transfected with NQO1 and flag tagged sirtuins (SIRT1-7) or p73b (as negative control in all the transfection experiments). Following immunoprecipitation with flag beads the association of NQO1 was analyzed by immunoblot with the indicated antibodies. B) Flag tagged SIRT1 or the SIRT1 H363Y mutant were co transfected in the presence of NQO1. The association of NQO1 was analyzed following immunoprecipitation with flag beads. C) Bacterially expressed and purified His-SIRT1 and NQO1 were incubated and the binding of NQO1 was analyzed following precipitation of His-Sirt1 with Ni-NTA beads. D) HEK293 over expressing flag tagged SIRT1 and NQO1 were treated with 10mM NAM or 50mM Resveratrol for 16 hours and the association of NQO1 was examined following immunoprecipitation with flag beads. E) Endogenous SIRT1 and NQO1 interact. Mouse liver extracts were immunoprecipitated (IP) with control empty beads, control IgG-coated beads, or anti-SIRT1-coated beads; and protein levels were analyzed by Western blotting (IB). The results of two anti-SIRT1 IPs are shown.

Nicotinamide (NAM), the byproduct of SIRT1 enzymatic activity, has been shown to inhibit SIRT1 activity whereas the natural polyphenol resveratrol has been shown to activate SIRT1 activity in cells (23). To examine whether SIRT1 activity modulators affect the binding to NQO1 we over expressed flag-SIRT1 with NQO1 in HEK293 cells and examined the co immunoprecipitation of NQO1 to flag-SIRT1 following treatment with either 10mM NAM or 50μM Resveratrol. These treatments did not affect NQO1 SIRT1 association (Fig. 1D). Next we used mouse liver extract to ask whether endogenous SIRT1 interacts with NQO1. SIRT1 was immunoprecipitated and the level of NQO1 was examined by immunoblotting with NQO1 specific antibody (Fig 1E). These data suggest that the over-expressed and the endogenous NQO1 directly interact with SIRT1.

### SIRT1 supports NQO1 expression

The binding of NQO1 to SIRT1 suggests there is a functional interplay between the two enzymes. Given the role of NQO1 in protein degradation we examined the possibility that SIRT1 accumulation is regulated by NQO1. To this end we used dicoumarol, a potent inhibitor of NQO1, which promotes the degradation of NQO1-protected proteins, such as p53 (18). SIRT1 protein levels were not affected by increased doses of dicoumarol that induced p53 protein degradation (Fig. 2A), Also NQO1 overexpression did not alter the level of the endogenous SIRT1 protein levels (Fig. 2B). Thus, NQO1 binding to SIRT1 does not induce functional stabilization of SIRT1 but rather might be due to other functional interplay.

**Figure 2.**
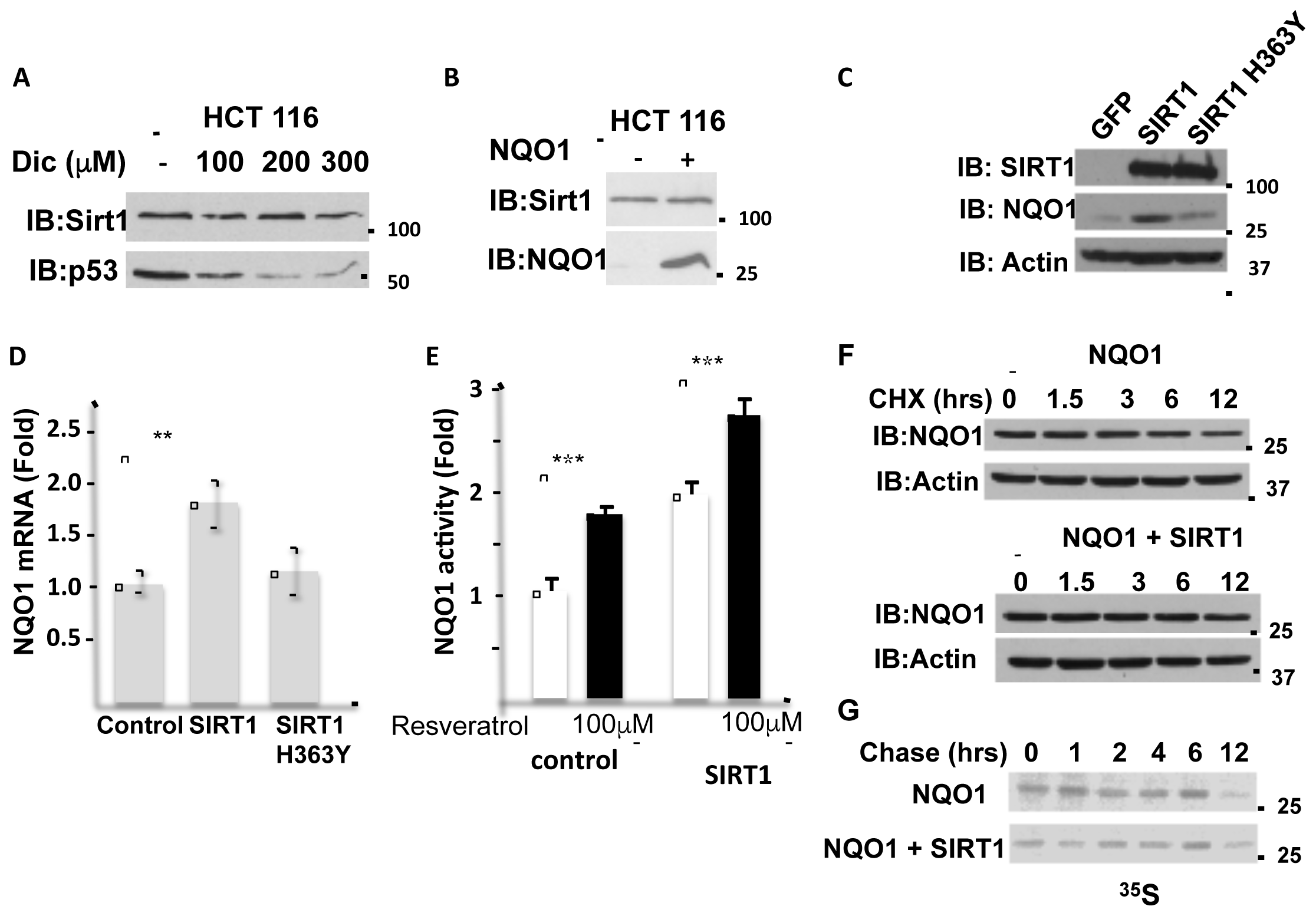
SIRT1 increases the endogenous NQO1 expression and protein level. A) HCT116 cells were treated with indicated concentrations of d¡coumarol and the level of SIRT1 and p53 were analyzed by Western blotting. B) The protein level of SIRT1 was analyzed in HCT116 control and NQO1 overexpressing cells.. NIH3T3 stably over expressing SIRT1, mutant H363Y SIRT1 or GFP as a control were examined for the NQO1 protein level (C), mRNA level (D) and activity (E) in the presence or absence of Resveratrol as indicated, for 16 hours. The stability of NQO1 was examined in HEK293 cells over expressing NQO1 in the presence or absence of flag-SIRT1 following cycloheximide treatment (100mg/ml) at indicated time points (F) or following pulse labeling with ^35^S Methionine and harvesting at indicated time points following the addition of unlabeled methionine (chase)(G).

Next we explored the effect of SIRT1 on NQO1 levels. To this end we infected NIH3T3 cells with a control GFP or with either wild type or H363Y mutant SIRT1 expressing plasmids and followed the level and the activity of the endogenous NQO1. SIRT1 over-expression in this system increased NQO1 protein levels compared to the control GFP or the mutant H363Y SIRT1 expressing cells (Fig. 2C). Similar results were obtained at the level of mRNA. In wild-type SIRT1 overexpressing cells, but not in the negative controls a 2-fold induction in NQO1 mRNA was observed (Fig. 2D). Next, we explored the NQO1 enzymatic activity. Control and SIRT1 over expressing cells were treated with Resveratrol for 16 hours. Following treatment, the cell extracts were analyzed for NQO1 enzymatic activity. Over expression of SIRT1 alone increased the level and the NQO1 activity by 2-fold whereas the addition of resveratrol further increased NQO1 activity by about 3-fold (Fig. 2E), possibly via an increase at the level of NQO1 expression (24).

Over expression of SIRT1 did not alter the half-life of NQO1 as analyzed by inhibiting translation with cycloheximide (Fig. 2F) or by ^35^S methionine pulse chase experiment (Fig. 2G). Our data suggest that active SIRT1 up regulated NQO1 expression.

### Mitochondrial inhibition induced NQO1 accumulation

NQO1 is an oxidoreductase that reduces various quinones by oxidizing NADH to NAD^+^. During mitochondrial inhibition there is an increase in the NADH/NAD^+^ ratio and ROS, and both are expected to induce NQO1 activity. To inhibit mitochondrial oxidative respiration, we used rotenone to inhibit complex-I, or antimycin A to inhibit complex-III (Fig 3A). These inhibitors increase the NADH/NAD^+^ ratio. NQO1 protein levels markedly increased following mitochondrial inhibition. This behavior was not cell type specific and observed in HCT 116, HEK293T and NIH3T3 cells (Fig. 3B-D).

**Figure 3.**
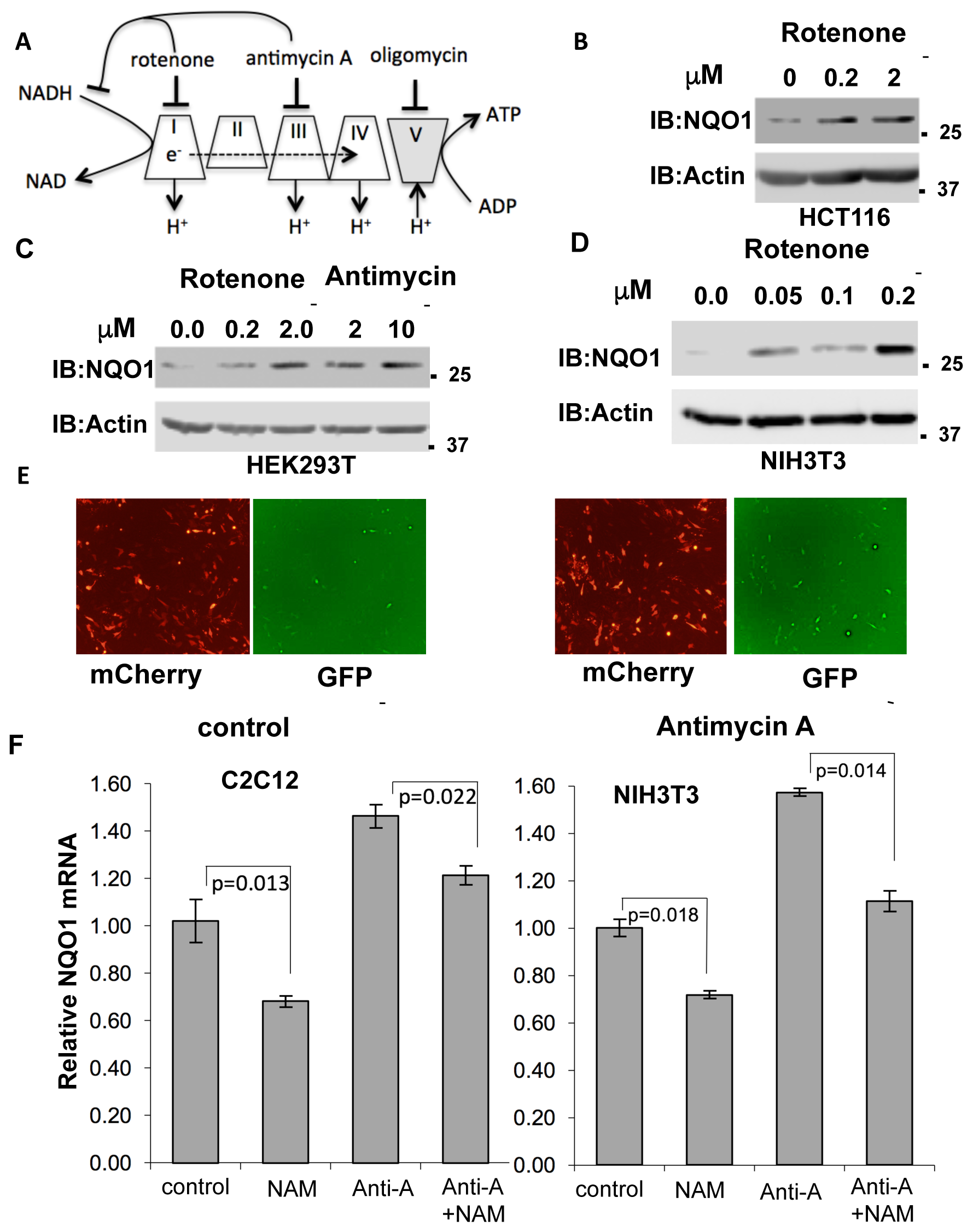
Mitochondrial inhibition induces NQO1 accumulation. A) Mitochondrial respiratory complexes (I-V) and their specific inhibitors are depicted. B-D). The endogenous levels of NQO1 protein were examined in HCT116 (B), HEK293T (C) and NIH3T3 (D) cells following treatment with indicated concentration of either rotenone (complex I inhibitor) or antimycin A (complex III inhibitor) for 16 hours. E) Using the pCDNA3 Peredox-mCherry reporter (25) we measured changes at the level of the NADH/NAD^+^ ratio (higher GFP/mCherry means higher NADH/NAD^+^ ratio). While the mCherry levels were not affected by antimycin A, GFP levels increased, indicating higher NADH levels. F) Inhibition of SIRT1 reduces NQO1 mRNA levels under basal and mitochondrial inhibition conditions. C2C12 or NIH3T3 cells were treated with antimycin A (10 mM) and NAM (10mM) as indicated for 16 h and analyzed for NQO1 mRNA expression by quantitative PCR.

Using the pCDNA3 Peredox-mCherry reporter (25) we show that the NADH/NAD^+^ ratio increases upon antimycin-A treatment as detected by higher GFP/mCherry ratio (Fig. 3E). Next we tested the NQO1 RNA level and found that it is increased by antimycin A treatment both in C2C12 and NIH3T3 cells (Fig. 3F). Interestingly, the NQO1 RNA levels are reduced by NAM, a known Sirtuins inhibitor. These results suggest that maximal induction of NQO1 expression under mitochondrial stress requires SIRT1 activity.

### Inhibition of the mitochondria increases nuclear NQO1 levels and NQO1-SIRT1 association

It has been reported that NAD^+^ related metabolic enzymes are involved in regulating transcription (12), and therefore we examined the cellular localization of NQO1. In untreated NIH3T3 cells the vast majority of the endogenous NQO1 was cytoplasmic (Fig. 4A). Under mitochondrial inhibition NQO1 could be detected in the nucleus (Fig. 4B). To further examine NQO1 nuclear localization under mitochondrial stress the endogenous NQO1 level in C2C12 cells was subjected to immunofluorescence analysis (Fig 4C). Antimycin A treatment markedly increased NQO1 nuclear localization. Similar results, although to a lesser extent were obtained with H1299 cells in which the endogenous NQO1 gene contains YFP in the second exon (clone 130207p11G9). Interestingly, under this setting the NQO1 nuclear localization was reduced by EX527, a SIRT1 specific inhibitor (Fig 4D).

**Figure 4.**
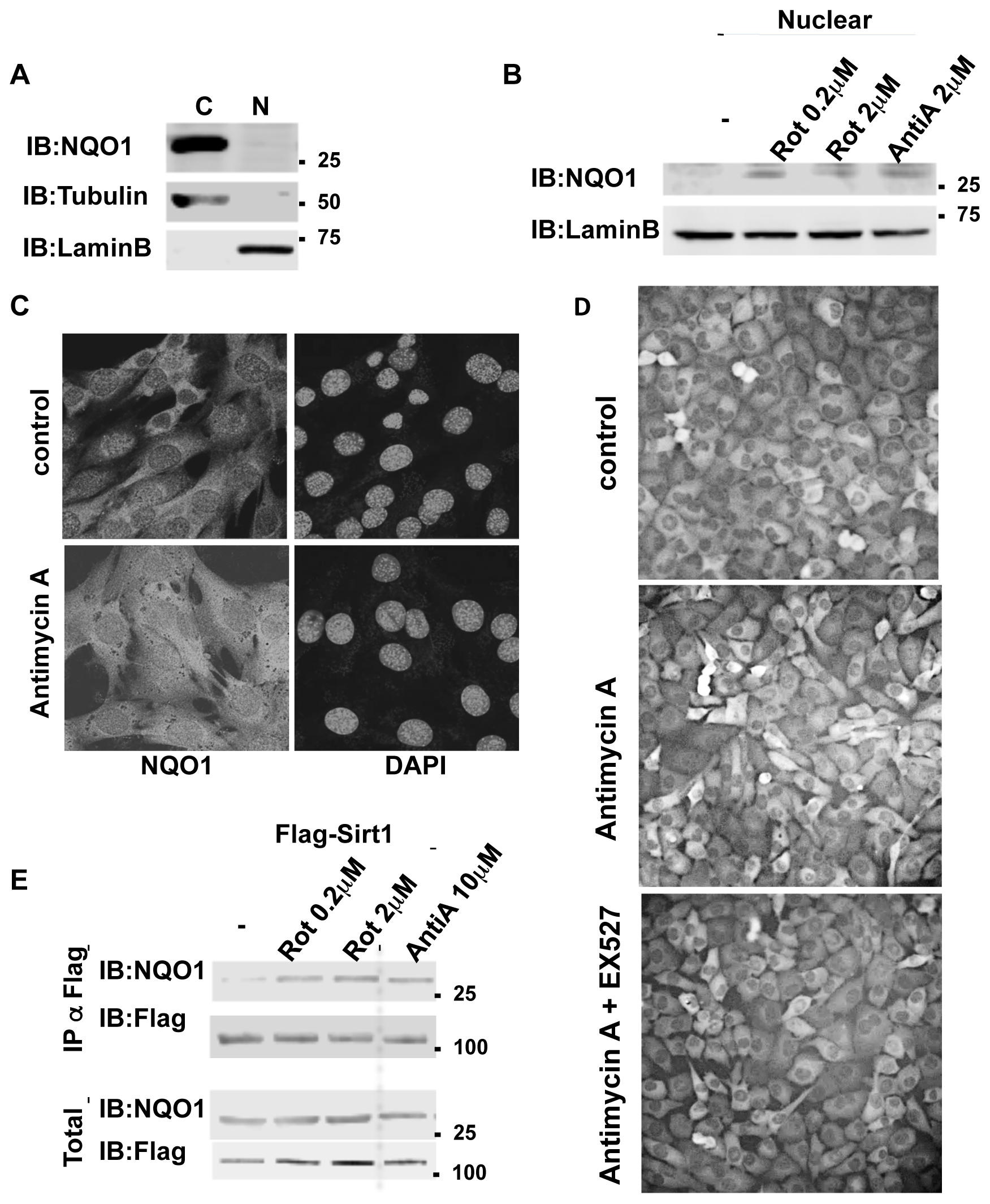
Mitochondrial inhibition increases nuclear NQO1 level and its SIRT1 association. A) cytoplasmatic (C) and nuclear (N) NQOldistribution were analyzed in NIH3T3 cells. B) The nuclear fraction of NQO1 was analyzed in NIH3T3 cells following treatment with indicated concentrations of rotenone or antimycin A for 16 hours. C) Subcellular distribution of NQO1 under conditions of mitochondrial inhibition. C2C12 cells were antimycin A treated (10 mM for 16 h) and fixed for immunofluorescence analysis with the NQO1 antibody (Epitomics 2618-1). Nuclear staining is indicated by DAPI. D) H1299 cells in which the endogenous NQO1 gene contains YFP in the second exon (clone 130207p11G9) were antimycin A treated in the presence or absence of EX527 (25 mM), a specific Sirt1 inhibitor. E) HEK293T cells over expressing Flag-SIRT1 and NQO1 were treated with indicated concentrations of rotenone or antimycin A for 16 hours. Following the treatment the nuclear fraction was isolated and the association of NQO1 to SIRT1 in the nucleus was analyzed by flag immunoprecipitation.

SIRT1 plays a protective role under mitochondrial inhibition conditions (26,27). This led us to explore the possibility that the NQO1-SIRT1 interaction is improved under these conditions. Flag-SIRT1 was immuno-precipitated (IP) and the coIP of NQO1 was analyzed in the control HEK293 cells and cells that were treated with different concentrations of Rotenone and antimycin A. NQO1 coIP with SIRT1 markedly increased with both mitochondrial inhibitors (Fig. 4E). These data suggest that during mitochondrial inhibition NQO1 is nuclear localized and that SIRT1-NQO1 physical interaction is increased.

### NQO1 supports SIRT1 regulated gene expression under mitochondrial inhibition

Next we wished to examine if NQO1 activity regulates SIRT1 function in transcription. We hypothesized that NQO1 stimulated SIRT1 deacetylase activity by converting NADH to the SIRT1 cofactor, NAD+. PGC-1α is a transcription coactivator involved in mitochondrial biogenesis (28) that is deacetylated by SIRT1, a process that potentiates PGC-1α activity. Interestingly, we have previously shown that NQO1 depletion increases PGC-1α acetylation (19) suggesting that NQO1 supports SIRT1 enzymatic function (Fig 5A).

**Figure 5.**
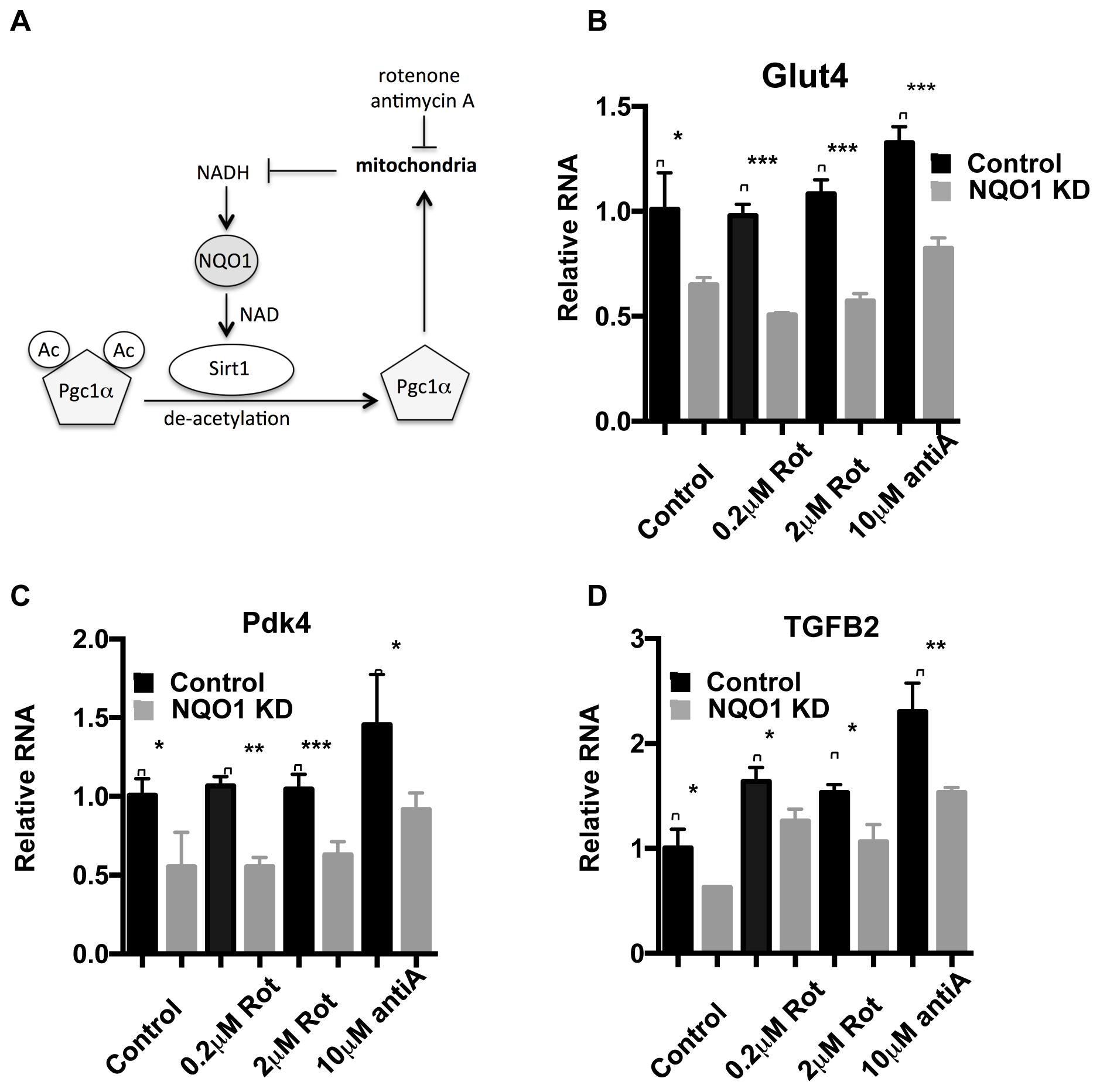
NQO1 supports SIRT1 regulated gene expression under mitochondrial inhibition. A) The interplay and the reported mechanism of NQO1-SIRT1 and PGC-la under mitochondrial stress are schematically presented. B-D) The mRNA levels of the SIRT1 regulated genes Glut4 (B) Pdk4 (C) and TGFB2 (D) were examined in NIH3T3 cells stably over expressing an NQO1 targeting shRNA or a control shRNA following treatment with indicated concentrations of rotenone (rot) or antimycin A (AntiA) for 16 hours. Statistical significance was analyzed by conducting two-tailed unpaired t-test. * p <0.05, ** p< 0.01, *** p<0.001

The SIRT1-PGC-1α circuit induces the expression of pyruvate dehydrogenase kinase-4 (PDK4) and Glucose transporter type 4 (GLUT4) (4). We next asked whether under mitochondrial stress NQO1 plays a role in regulating these genes. To this end we depleted NQO1 by the knockdown strategy (19). Interestingly, we found that under NQO1 depletion the expression of these genes was compromised (Fig. 5B-C).

Nicotinamide phosphoribosyltransferase (NAMPT) and nicotinamide mononucleotide adenylyltransferase 1 (NMNAT-1) salvage pathway regulates NAD level and SIRT1 activity. One of the cellular genes highly affected by depletion of any of these enzymes is the transforming growth factor beta 2 (TGFB2) gene (12). We examined the role of NQO1 in regulating SIRT1 mediated transcription of TGFB2. Similar to the role of NQO1 in the SIRT1-PGC-1α circuit, we found that NQO1 regulates the SIRT1-NAD salvage circuit as well (Fig 5D). These results suggest that NQO1 potentiates the transcription of the SIRT1 target genes in a more systemic fashion.

### NQO1-SIRT1 circuit determines cell fate

Having revealed the NQO1-SIRT1 circuit in regulating gene expression under mitochondrial stresses we next investigated the role of this circuit in cell fate determination. It has been reported that nuclear SIRT1 protects C2C12 cells from antimycin A-induced cell death (26). We asked whether NQO1 regulates the SIRT1-mediated protective role. We found that NQO1-depleted C2C12 cells are less viable under antimycin-A treatment (Fig 6A), whereas over-expression of SIRT1 increased cellular viability as previously described. Remarkably, the role of SIRT1 in maintaining viability is markedly blunted in NQO1-depleted cells. These data suggest that under mitochondrial stress NQO1 positively regulates nuclear SIRT1 in cell fate determination (Fig 6B).

**Figure 6.**
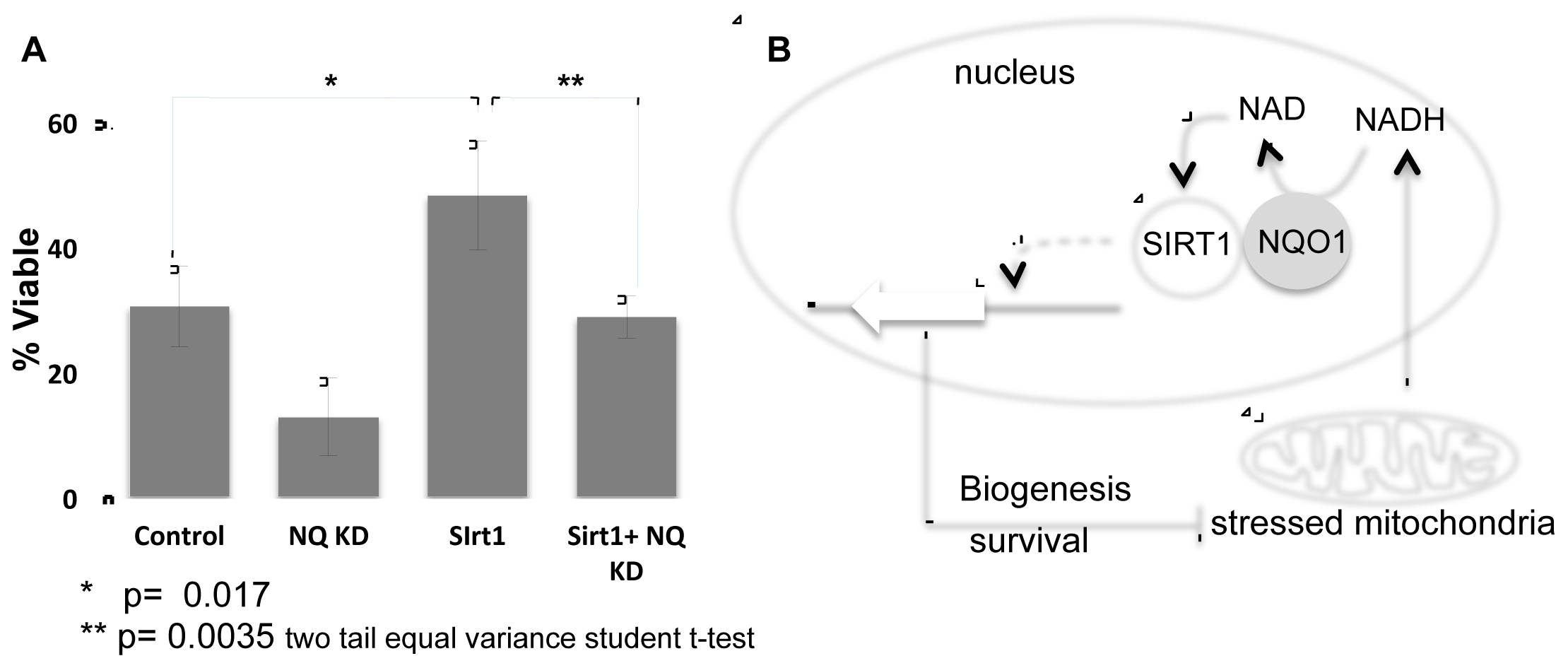
NQO1-SIRT1 circuit determines cell fate. A) C2C12 cells either over expressing SIRT1 or the shRNA for NQO1 or both were treated with Antimycin A and the viability was examined after 48 hours. B) The NADH metabolism in regulating the SIRT1-NQO1-PGC-1a feedback loop and mitochondrial biogenesis are schematically summarized.

## DISCUSSION

In this work we functionally coupled NQO1 with SIRT1. We show that NQO1 physically interacts with the enzymatically active SIRT1. The NADH/NAD^+^ ratio in the cells is a rheostatic process whose set-point is dictated mainly by the mitochondria. NQO1 activity increases NAD^+^, the SIRT1 cofactor, coupling these enzymes in a functional module. We also provide evidence for a positive feed-forward loop whereby SIRT1 up-regulates NQO1 level. The observed interplay between NQO1 and SIRT1 establishes a new regulatory circuit sensing the NADH/NAD^+^ ratio. This circuitry is important in particular under conditions where the NADH/NAD^+^ levels are high such as mitochondrial stress. Under NQO1 depletion the ability of SIRT1 to maintain cell viability following mitochondrial inhibition is compromised. Our work suggests that the NQO1-SIRT1 regulatory circuit is an important component in cell fate decision-making following mitochondrial stress.

The regulatory circuit under study is activated by mitochondrial stress, conditions whereby PGC-1α is activated. Recently we have shown that NQO1 increases PGC-1α protein level in an NADH-dependent manner. The accumulated PGC-1α induces the expression of metabolic and mitochondrial biogenesis genes (19). We have now expanded this circuit by demonstrating that not only PGC-1α but also SIRT1 is regulated by NQO1. The ability of NQO1 to alter the NADH/NAD^+^ ratio attributes a central role to NQO1 in this regulatory PGC-1α/-SIRT1-NQO1 module.

The role of NQO1 described here adds another layer in the moonlighting cellular function of NQO1. NQO1 functions as a quinone reductase, one of the cellular phase-two detoxifying enzymes (13). However, NQO1 is also involved in transcription regulation, translation and protein degradation (29-31). NQO1 is associated with the 20S proteasome (31) and regulates the stability of numerous intrinsically disordered proteins (IDPs) such as p53, p73, and PGC1α in an NADH-dependent manner (19,31). This moonlighting behavior of a “metabolic enzyme” is not a unique property of NQO1. The PKM2 and GAPDH metabolic proteins are involved in numerous cellular processes. GAPDH for instance was shown to be involved in transcription regulation (32) and as a chaperone of newly synthesized ribosomal protein L13a (33). To some extent these GAPDH activities are also NADH-dependent.

SIRT1 functions have been highly explored in the context of cellular aging and increased yeast life span (34). In the mouse model SIRT1 activity is beneficial in preventing pathologies associated with aging in tissues like muscle, brain and heart (2). Although SIRT1 protein levels are high in some aged tissues its activity however remains highly dependent on the NAD+/NADH ratio (20,30). We have recently shown that NQO1 protein levels significantly increase in the aging brain (20). Moreover, increasing the NAD+/NADH ratio by facilitation of NQO1 activity is sufficient to delay muscle and brain functional decline in mice (35). These processes have been described to involve SIRT1, attributing an important role to the NQO1-SIRT1 circuit in aging.

In yeast the NQO1 homologue NQR1 directly increases both chronological and replicative lifespan. The replicative life span extension is induced by shifting cells towards respiratory metabolism, a process mediated by the SIRT1 yeast homologue SIR2 (36). Thus, the functional link between the NADH oxidoreductases NQO1 (NQR1) and SIRT1 (SIR2) is evolutionarily conserved linking cellular metabolism to epigenetic regulation.

NADH/NAD^+^ levels and ratio are emerging as a critical cellular redox/metabolic sensing ratio regulating epigenetic, transcription, protein degradation and cellular metabolism (37). In the NQO1 knockout mouse one of the most significant effects reported is the alteration in the NADH/NAD^+^ ratio (38), signifying the role of NQO1 in maintaining the cellular NADH/NAD^+^ levels. NQO1 ability to regulate the cellular NADH/NAD^+^ ratio is a key component of the plasma membrane redox system (PMRS) as well and highly affects cellular viability during oxidative and metabolic stresses (39). However, the mechanism by which NQO1 mediates these functions is not yet well understood. We believe that the crosstalk between SIRT1 and NQO1 could directly link the NQO1 redox function with the cellular regulatory one.

NAD^+^ hydrolysis is an essential step in the SIRT1 de-acetylation reaction. One molecule of NAD^+^ is consumed and one molecule of o-acetyl-ADP ribose and nicotinamide is produced for every lysine de-acetylation reaction (1). This NAD^+^ dependence links cellular metabolism with SIRT1 function in cells and suggests that physiological conditions that alter intracellular NAD^+^ levels or metabolic enzymes involved in NADH/NAD^+^ metabolism may regulate SIRT1 deacetylase activity, and the biological outcome. This has been shown to be true in yeast where mutation of the NPT1 gene encoding a nicotinate phosphoribosyltransferase, a critical enzyme in the NAD^+^ salvage pathway reduces NAD^+^ level and inhibits Sir2 mediated silencing (40). In mammalian cells the NAD^+^ salvage pathway enzymes NAMNT-1 and NAMPT both increase NAD^+^ levels and activate SIRT1-mediated transcription regulation of a number of genes, including TGFB2 (41). We show here that at least in the case of TGFB2 its transcription activation by SIRT1 is partially compromised under NQO1 depletion. Interestingly, PGC-1α itself has been reported to induce NAD biosynthesis (42). NAD has a central role in energy metabolism and is rate-limiting for mitochondrial function (10,43). Therefore, NQO1, SIRT1 and PGC-1α are the core components of a metabolic rheostatic module responding to mitochondrial stress by sensing the NAD^+^/NADH ratio.

## Acknowledgments

We would like to acknowledge Eric Verdin, Robert Weinberg and Haim Cohen for the different plasmids and Eyal Kimchi for his assistance. This work was supported by the Israel Science Foundation, Grant no. 1591/15.

The authors declare that they have no conflicts of interest with the contents of this article.

## References

1. Blander, G., and Guarente, L. (2004) The Sir2 family of protein deacetylases. Annu Rev Biochem 73, 417–435

2. Haigis, M. C., and Sinclair, D. A. (2010) Mammalian sirtuins: biological insights and disease relevance. Annu Rev Pathol 5, 253–295

3. Nemoto, S., Fergusson, M. M., and Finkel, T. (2005) SIRT1 functionally interacts with the metabolic regulator and transcriptional coactivator PGC-1{alpha}. J Biol Chem 280, 16456–16460

4. Canto, C., Gerhart-Hines, Z., Feige, J. N., Lagouge, M., Noriega, L., Milne, J. C., Elliott, P. J., Puigserver, P., and Auwerx, J. (2009) AMPK regulates energy expenditure by modulating NAD+ metabolism and SIRT1 activity. Nature 458, 1056–1060

5. Kaeberlein, M., McVey, M., and Guarente, L. (1999) The SIR2/3/4 complex and SIR2 alone promote longevity in Saccharomyces cerevisiae by two different mechanisms. Genes Dev 13, 2570–2580

6. Asher, G., Gatfield, D., Stratmann, M., Reinke, H., Dibner, C., Kreppel, F., Mostoslavsky, R., Alt, F. W., and Schibler, U. (2008) SIRT1 regulates circadian clock gene expression through PER2 deacetylation. Cell 134, 317–328

7. Fulco, M., Schiltz, R. L., Iezzi, S., King, M. T., Zhao, P., Kashiwaya, Y., Hoffman, E., Veech, R. L., and Sartorelli, V. (2003) Sir2 regulates skeletal muscle differentiation as a potential sensor of the redox state. Mol Cell 12, 51–62

8. Canto, C., Jiang, L. Q., Deshmukh, A. S., Mataki, C., Coste, A., Lagouge, M., Zierath, J. R., and Auwerx, J. (2010) Interdependence of AMPK and SIRT1 for metabolic adaptation to fasting and exercise in skeletal muscle. Cell metabolism 11, 213–219

9. Hayashida, S., Arimoto, A., Kuramoto, Y., Kozako, T., Honda, S., Shimeno, H., and Soeda, S. (2010) Fasting promotes the expression of SIRT1, an NAD+ - dependent protein deacetylase, via activation of PPARalpha in mice. Molecular and cellular biochemistry 339, 285–292

10. Bai, P., Canto, C., Oudart, H., Brunyanszki, A., Cen, Y., Thomas, C., Yamamoto, H., Huber, A., Kiss, B., Houtkooper, R. H., Schoonjans, K., Schreiber, V., Sauve, A. A., Menissier-de Murcia, J., and Auwerx, J. (2011) PARP-1 inhibition increases mitochondrial metabolism through SIRT1 activation. Cell metabolism 13, 461–468

11. Canto, C., Houtkooper, R. H., Pirinen, E., Youn, D. Y., Oosterveer, M. H., Cen, Y., Fernandez-Marcos, P. J., Yamamoto, H., Andreux, P. A., Cettour-Rose, P., Gademann, K., Rinsch, C., Schoonjans, K., Sauve, A. A., and Auwerx, J. (2012) The NAD(+) precursor nicotinamide riboside enhances oxidative metabolism and protects against high-fat diet-induced obesity. Cell metabolism 15, 838–847

12. Zhang, T., Berrocal, J. G., Frizzell, K. M., Gamble, M. J., DuMond, M. E., Krishnakumar, R., Yang, T., Sauve, A. A., and Kraus, W. L. (2009) Enzymes in the NAD+ salvage pathway regulate SIRT1 activity at target gene promoters. J Biol Chem 284, 20408–20417

13. Lind, C., Cadenas, E., Hochstein, P., and Ernster, L. (1990) DT-diaphorase: purification, properties, and function. Methods Enzymol 186, 287–301

14. Uversky, V. N., and Longhi, S. (2010) Instrumental analysis of intrinsically disordered proteins: assessing structure and conformation, Wiley, Hoboken, N.J.

15. Talalay, P., and Dinkova-Kostova, A. T. (2004) Role of nicotinamide quinone oxidoreductase 1 (NQO1) in protection against toxicity of electrophiles and reactive oxygen intermediates. Methods Enzymol 382, 355–364

16. Gaikwad, A., Long, D. J., 2nd, Stringer, J. L., and Jaiswal, A. K. (2001) In vivo role of NAD(P)H:quinone oxidoreductase 1 (NQO1) in the regulation of intracellular redox state and accumulation of abdominal adipose tissue. J Biol Chem 276, 22559–22564

17. Haefeli, R. H., Erb, M., Gemperli, A. C., Robay, D., Courdier Fruh, I., Anklin, C., Dallmann, R., and Gueven, N. (2011) NQO1-dependent redox cycling of idebenone: effects on cellular redox potential and energy levels. PLoS One 6, e17963

18. Asher, G., Lotem, J., Cohen, B., Sachs, L., and Shaul, Y. (2001) Regulation of p53 stability and p53-dependent apoptosis by NADH quinone oxidoreductase 1. Proceedings of the National Academy of Sciences of the United States of America 98, 1188–1193

19. Adamovich, Y., Shlomai, A., Tsvetkov, P., Umansky, K. B., Reuven, N., Estall, J. L., Spiegelman, B. M., and Shaul, Y. (2013) The Protein Level of PGC-1alpha, a Key Metabolic Regulator, Is Controlled by NADH-NQO1. Mol Cell Biol 33, 2603–2613

20. Tsvetkov, P., Adamovich, Y., Elliott, E., and Shaul, Y. (2011) E3 ligase STUB1/CHIP regulates NAD(P)H:quinone oxidoreductase 1 (NQO1) accumulation in aged brain, a process impaired in certain Alzheimer disease patients. The Journal of biological chemistry 286, 8839–8845

21. Prochaska, H. J., and Santamaria, A. B. (1988) Direct measurement of NAD(P)H:quinone reductase from cells cultured in microtiter wells: a screening assay for anticarcinogenic enzyme inducers. Anal Biochem 169, 328–336

22. Tsvetkov, P., Reuven, N., Prives, C., and Shaul, Y. (2009) Susceptibility of p53 unstructured N terminus to 20 S proteasomal degradation programs the stress response. J Biol Chem 284, 26234–26242

23. Howitz, K. T., Bitterman, K. J., Cohen, H. Y., Lamming, D. W., Lavu, S., Wood, J. G., Zipkin, R. E., Chung, P., Kisielewski, A., Zhang, L. L., Scherer, B., and Sinclair, D. A. (2003) Small molecule activators of sirtuins extend Saccharomyces cerevisiae lifespan. Nature 425, 191–196

24. Hsieh, T. C., Lu, X., Wang, Z., and Wu, J. M. (2006) Induction of quinone reductase NQO1 by resveratrol in human K562 cells involves the antioxidant response element ARE and is accompanied by nuclear translocation of transcription factor Nrf2. Med Chem 2, 275–285

25. Hung, Y. P., Albeck, J. G., Tantama, M., and Yellen, G. (2011) Imaging cytosolic NADH-NAD(+) redox state with a genetically encoded fluorescent biosensor. Cell metabolism 14, 545–554

26. Tanno, M., Sakamoto, J., Miura, T., Shimamoto, K., and Horio, Y. (2007) Nucleocytoplasmic shuttling of the NAD+-dependent histone deacetylase SIRT1. J Biol Chem 282, 6823–6832

27. Gerhart-Hines, Z., Rodgers, J. T., Bare, O., Lerin, C., Kim, S. H., Mostoslavsky, R., Alt, F. W., Wu, Z., and Puigserver, P. (2007) Metabolic control of muscle mitochondrial function and fatty acid oxidation through SIRT1/PGC-1alpha. The EMBO journal 26, 1913–1923

28. Wu, Z., Puigserver, P., Andersson, U., Zhang, C., Adelmant, G., Mootha, V., Troy, A., Cinti, S., Lowell, B., Scarpulla, R. C., and Spiegelman, B. M. (1999) Mechanisms controlling mitochondrial biogenesis and respiration through the thermogenic coactivator PGC-1. Cell 98, 115–124

29. Adler, J., Reuven, N., Kahana, C., and Shaul, Y. (2010) c-Fos proteasomal degradation is activated by a default mechanism, and its regulation by NAD(P)H:quinone oxidoreductase 1 determines c-Fos serum response kinetics. Mol Cell Biol 30, 3767–3778

30. Alard, A., Fabre, B., Anesia, R., Marboeuf, C., Pierre, P., Susini, C., Bousquet, C., and Pyronnet, S. (2010) NAD(P)H quinone-oxydoreductase 1 protects eukaryotic translation initiation factor 4GI from degradation by the proteasome. Mol Cell Biol 30, 1097–1105

31. Asher, G., Tsvetkov, P., Kahana, C., and Shaul, Y. (2005) A mechanism of ubiquitin-independent proteasomal degradation of the tumor suppressors p53 and p73. Genes Dev 19, 316–321

32. Zheng, L., Roeder, R. G., and Luo, Y. (2003) S phase activation of the histone H2B promoter by OCA-S, a coactivator complex that contains GAPDH as a key component. Cell 114, 255–266

33. Jia, J., Arif, A., Willard, B., Smith, J. D., Stuehr, D. J., Hazen, S. L., and Fox, P. L. (2012) Protection of extraribosomal RPL13a by GAPDH and dysregulation by S-nitrosylation. Molecular cell 47, 656–663

34. Kaeberlein, M., McVey, M., and Guarente, L. (1999) The SIR2/3/4 complex and SIR2 alone promote longevity in Saccharomyces cerevisiae by two different mechanisms. Genes & development 13, 2570–2580

35. Lee, J. S., Park, A. H., Lee, S. H., Kim, J. H., Yang, S. J., Yeom, Y. I., Kwak, T. H., Lee, D., Lee, S. J., Lee, C. H., Kim, J. M., and Kim, D. (2012) Beta-Lapachone, a Modulator of NAD Metabolism, Prevents Health Declines in Aged Mice. PLoS One 7, e47122

36. Jimenez-Hidalgo, M., Santos-Ocana, C., Padilla, S., Villalba, J. M., Lopez-Lluch, G., Martin-Montalvo, A., Minor, R. K., Sinclair, D. A., de Cabo, R., and Navas, P. (2009) NQR1 controls lifespan by regulating the promotion of respiratory metabolism in yeast. Aging Cell 8, 140–151

37. Ying, W. (2008) NAD+/NADH and NADP+/NADPH in cellular functions and cell death: regulation and biological consequences. Antioxid Redox Signal 10, 179–206

38. Gaikwad, A., Long, D. J., 2nd, Stringer, J. L., and Jaiswal, A. K. (2001) In vivo role of NAD(P)H:quinone oxidoreductase 1 (NQO1) in the regulation of intracellular redox state and accumulation of abdominal adipose tissue. The Journal of biological chemistry 276, 22559–22564

39. Tan, A. S., and Berridge, M. V. (2010) Evidence for NAD(P)H:quinone oxidoreductase 1 (NQO1)-mediated quinone-dependent redox cycling via plasma membrane electron transport: A sensitive cellular assay for NQO1. Free Radic Biol Med 48, 421–429

40. Sandmeier, J. J., Celic, I., Boeke, J. D., and Smith, J. S. (2002) Telomeric and rDNA silencing in Saccharomyces cerevisiae are dependent on a nuclear NAD(+) salvage pathway. Genetics 160, 877–889

41. Zhang, T., Berrocal, J. G., Frizzell, K. M., Gamble, M. J., DuMond, M. E., Krishnakumar, R., Yang, T., Sauve, A. A., and Kraus, W. L. (2009) Enzymes in the NAD+ salvage pathway regulate SIRT1 activity at target gene promoters. The Journal of biological chemistry 284, 20408–20417

42. Tran, M. T., Zsengeller, Z. K., Berg, A. H., Khankin, E. V., Bhasin, M. K., Kim, W., Clish, C. B., Stillman, I. E., Karumanchi, S. A., Rhee, E. P., and Parikh, S. M. (2016) PGC1alpha drives NAD biosynthesis linking oxidative metabolism to renal protection. Nature 531, 528–532

43. Gomes, A. P., Price, N. L., Ling, A. J., Moslehi, J. J., Montgomery, M. K., Rajman, L., White, J. P., Teodoro, J. S., Wrann, C. D., Hubbard, B. P., Mercken, E. M., Palmeira, C. M., de Cabo, R., Rolo, A. P., Turner, N., Bell, E. L., and Sinclair, D. A. (2013) Declining NAD(+) induces a pseudohypoxic state disrupting nuclear-mitochondrial communication during aging. Cell 155, 1624–1638

